# Dairy DigiD ∼ A Deep Learning-Based, Non-Invasive Biometric Identification System for Dairy Cattle Using Detectron2

**DOI:** 10.1101/2024.12.14.628477

**Authors:** Shubhangi Mahato, Hanqing Bi, Suresh Neethirajan

## Abstract

Precision livestock farming demands accurate and reliable individual animal identification for optimizing health monitoring, resource allocation, and overall herd productivity. Conventional identification methods—such as ear tagging and wearable sensors—are often invasive, potentially causing animal discomfort and affecting natural behaviors. To address these challenges, we introduce Dairy DigiD, a non-invasive biometric classification system for dairy cattle using facial image analysis. Leveraging deep learning and the Detectron2 framework, Dairy DigiD classifies cattle into four categories—young (<2 years), mature milking, pregnant, and old—using over 2,500 high-resolution facial images. A DenseNet121 model achieved a classification accuracy of 97%, showcasing its strong discriminative capability. Further, Detectron2 attained an average recall of 87% in category identification, an average precision of 96% in cattle detection, and an overall detection accuracy of 93%, demonstrating its robustness and adaptability to varied environmental conditions. Unlike traditional approaches that rely on fixed facial landmarks and can be sensitive to environmental variability, Dairy DigiD’s flexible deep learning architecture enables stable and scalable facial feature extraction. This non-invasive method minimizes animal stress, enhances data integrity, and facilitates improved herd management and welfare monitoring. In comparison to existing convolutional neural network (CNN)-based strategies, Dairy DigiD exhibits superior adaptability to real-world farm conditions, including variable lighting and complex poses. This work highlights the potential of integrating advanced computer vision and deep learning methodologies into precision dairy farming, offering an ethically aligned, efficient, and scalable approach to modern livestock management.

## 1. Introduction

Digital livestock farming has emerged as a critical approach to meet the growing global demand for sustainable and efficient agricultural practices (Neethirajan, 2021). Central to this paradigm is the need for accurate, reliable, and ethically sound methods of individual animal identification. This foundational capability underpins a wide range of precision management techniques, from optimized resource allocation to enhanced health monitoring and improved welfare. Historically, cattle identification relied on physical markers like ear tags and branding, overseen by bodies such as the Canadian Food Inspection Agency (CFIA). While long standard, these methods present inherent drawbacks. Physical identifiers may induce stress or discomfort, potentially altering natural behaviors. Even less invasive wearable devices such as collars or accelerometers can disrupt normal routines, raising questions about data integrity and animal well-being.

In pursuit of more refined, animal-centric techniques, researchers explored biometric methods like muzzle printing and coat pattern analysis (Hossain et al., (2022)l Li et al., (2022); Chen et al., (2021); Manoj et al., (2021)). Early computer vision efforts using Scale-Invariant Feature Transform (SIFT) and Random Sample Consensus (RANSAC) attempted to identify cattle through unique facial traits akin to human fingerprints. Barrera et al. (2013) demonstrated this potential with muzzle prints. Yet these early approaches struggled with scaling in real-world farms, hampered by variable lighting, cluttered backgrounds, and diverse poses. Strict reliance on precise image registration proved fragile, limiting large-scale deployment.

Subsequent machine learning methods automated feature extraction and classification, employing biometric traits with Support Vector Machines (SVM), k-Nearest Neighbors (k-NN), and Artificial Neural Networks (ANN). These approaches surpassed manual monitoring, easing workloads and improving accuracy. Still, they required hand-crafted features, restricting adaptability and robust performance in heterogeneous contexts. Small datasets further constrained generalization, diminishing their effectiveness in large-scale farming.

The advent of deep learning, especially Convolutional Neural Networks (CNNs), transformed cattle identification by automatically extracting and classifying features with greater precision. CNN-based architectures (e.g., ResNet, YOLO) delivered remarkable gains in image classification. El-Henawy et al. (2016) and Andrew et al. (2017) reported substantial improvements using CNNs combined with texture analysis and R-CNN methods on Friesian cattle. By 2021, CNN-based models like ResNet and YOLO neared 98.99% accuracy, enabling practical field-level implementations that bolster herd management, disease prevention, and breeding programs.

Despite these advances, current deep learning models still face persistent hurdles. Architectures like VGG16, ResNet, and YOLO excel in some scenarios but struggle in others. VGG16 achieves strong accuracy under controlled conditions but requires significant computation and lengthy training. YOLO excels in real-time localization within dynamic settings but may generate false positives for small objects like cow heads if not meticulously tuned. While YOLO and Faster RCNN excel at detection, they can lag behind VGG16 in classification speed. Nonetheless, YOLO’s versatility, spanning tasks from object detection to image captioning, enhances its adaptability.

These ongoing challenges underscore the need for class-agnostic deep learning frameworks that surpass existing architectural boundaries, particularly amid unpredictable farm conditions with uneven lighting, background noise, and complex cow poses. Environmental variability can undermine current models’ accuracy, as Neethirajan (2023) noted. Evaluations of suitable architectures naturally consider DenseNet121 (Huang et al., 2017), known for effective feature propagation and mitigated vanishing gradients. Preliminary tests on cow images hinted at robust recognition but fell short under challenging lighting, poses, and clutter, indicating the need to preserve DenseNet’s advantages while remedying its weaknesses.

Armed with these insights, we turned to Detectron2, a state-of-the-art object detection framework developed by Facebook AI Research. Detectron2 surpasses existing models in adaptability, precision, and accuracy under diverse farm environments. Its modular design and key-point detection capabilities assist in pinpointing subtle cattle features. Insights from DenseNet guided Detectron2’s pipeline, emphasizing the efficient extraction of subtle yet discriminative elements. Detectron2’s modularity supports custom pipelines for specialized uses like cattle identification. Built on PyTorch and CUDA, it delivers speed, accuracy, and adaptability, supporting real-time detection and reliable classification. Its 30-point keypoint detection enables more granular, stable assessments than standard algorithms, and its ability to fine-tune existing models expands its role in precision livestock management.

Prior cattle identification work often focused on muzzle patterns, ear tags, or basic CNNs, but Detectron2’s potential remains largely unexplored. By merging cutting-edge computer vision with deep learning, our proposed Dairy DigiD system aims for stable, accurate cattle identification in dynamic farm scenarios. This synergy promises enhanced scalability, accuracy, and minimal animal stress, aiding farmers in improved herd tracing, targeted data collection, and comprehensive welfare monitoring. Following a defined workflow (Figure 1), we begin by ensuring diverse, complex data. Preprocessing refines image quality, and annotating key features via the Computer Vision Annotation Tool (CVAT) informs model configuration and hyperparameter tuning. Ultimately, training and evaluating within Detectron2 produces an optimized identification model.

**Figure 1.**
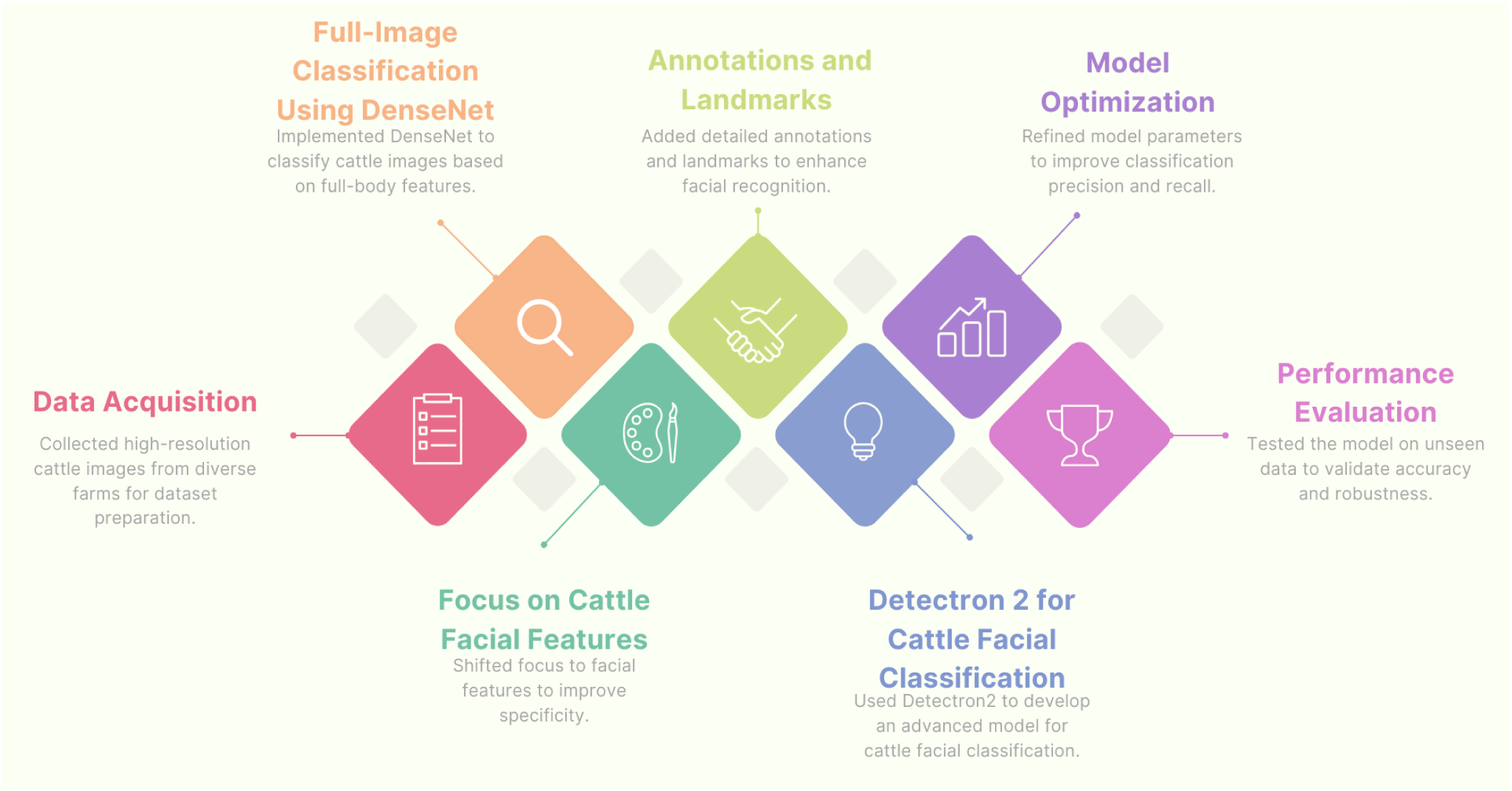
Workflow diagram illustrating the key steps in developing the Dairy DigiD cattle image classification model, from data collection to model evaluation.

This study addresses essential questions: (1) Can deep learning models like Detectron2 enhance cattle identification amid varying farm conditions? (2) How does Detectron2 compare to pre-trained models like VGG16 or GAN-based (VIA-masks) regarding accuracy, speed, and feasibility? We benchmark Detectron2 against established models, examining precision, recall, and F1-scores across categories—young, mature milking, pregnant, and old cows. Multiple deep learning solutions, including VGG, ResNet, and YOLO, have aided cattle recognition [22][28]. VGG16 excels in stable settings, YOLO thrives in dynamic localization but may misclassify small targets, and YOLO/Faster RCNN might trail VGG16 in classification speed. Yet YOLO’s adaptability remains a key asset. Such constraints highlight the need for class-agnostic deep learning in noisy, unstable conditions [19]. We posit Detectron2 outperforms existing models in flexibility, accuracy, and resilience. Leveraging Detectron2, a premier framework from Facebook AI Research, we tackle these challenges. Its customizable pipelines support real-time detection. Its 30-point keypoint detection surpasses traditional algorithms, enabling richer classification. Detectron2 also eases model creation and refinement, ideally suited for precision livestock tasks. While past studies explored muzzle patterns, ear tags, or CNN-based approaches, Detectron2 remains underexploited. By integrating advanced vision techniques with deep learning, Dairy DigiD ensures consistent, resilient identification as conditions shift. This guarantees scalability, non-invasive data collection, and high accuracy, benefiting farmers through enhanced herd management, data-driven decisions, and better welfare with minimal impact. Our work builds on prior advances but acknowledges practical limits. Machine learning once required manual features and struggled to scale. Deep learning improved automation and accuracy but environmental sensitivities and computational burdens persist. Detectron2’s modular, keypoint-based approach offers a promising path, adapting to intricate farm conditions. Emerging methods, like Vision Transformers (ViTs) combined with CNNs, transcend classical constraints. ViTs’ self-attention captures multi-scale features under tough conditions. Zhang et al. (2024) show ViT-based models can outperform CNNs in challenging scenarios. Integrating ViTs with Detectron2 may yield more accurate, scalable solutions for precision livestock management.

Our main objective is to evaluate Detectron2’s effectiveness in boosting cattle identification accuracy via Dairy DigiD, advancing precision livestock management. Specific aims include: i) Assessing Detectron2’s accuracy, speed, and deployability under varied conditions—uneven lighting, complex backdrops, diverse poses. ii) Comparing Detectron2’s performance with other pre-trained models (e.g., VGG16, VIA-masks) on accuracy, speed, and scalability to identify strengths and improvement areas. iii) Developing a scalable, non-invasive identification method that reduces animal stress, refines livestock management, improves disease control, and safeguards welfare, ensuring feasibility in large-scale farming.

## 2. Materials and Methods

The dataset used in this study is a carefully curated compilation designed to represent the diversity and complexity of dairy cattle populations in Nova Scotia, Atlantic Canada. It comprises more than 2,500 images of 180 cows from two breeds, forming a robust foundation for training advanced AI models capable of accurately identifying and analyzing individual animals.

### 2.1 Ethical Approvals

Prior to initiating data collection, all requisite approvals were secured from the university and the participating farm owners. The university’s ethics committee thoroughly examined the project, confirming adherence to guidelines and verifying that no invasive procedures were involved— only image and video capture of the cattle. Farm owners were fully briefed on the study’s objectives and methodologies, and their informed consent was obtained, ensuring openness and collaboration. Since no physical contact with the cattle occurred, no additional approvals were necessary beyond these initial clearances.

### 2.2 Cattle Demographics and Dataset Composition

Images were sourced from local dairy farms in Nova Scotia, with herd sizes varying from 60 to 110 animals. The cows ranged freely, producing images with backgrounds featuring barns, trees, and other cattle—crucial for developing AI models resilient to differing farm environments. Additional images were taken at Dalhousie University’s Agriculture Campus Ruminant Animal Centre, where cattle were briefly tethered. The dataset includes Holstein and Jersey breeds, focusing primarily on Holstein, and encompasses various ages and physiological states:

> Young cows (<2 years): 266 images
>
> Mature milking cows: 1315 images
>
> Pregnant cows: 89 images
>
> Old cows: 625 images

### 2.3 Data Collection Methods

High-resolution images and K videos were obtained using devices like the Samsung S21 (64MP, f/1.8, 26mm, PDAF, OIS), iPhone 14 (48MP, f/1.78, 24mm, sensor-shift OIS), and iPhone 13 (12MP, f/1.6, 26mm, dual pixel PDAF, sensor-shift OIS). This variety ensured distinct image qualities, facilitating robust data that captures details under different lighting conditions and angles. Employing multiple devices and resolutions improved color accuracy and flexibility. Ultimately, more than 3,000 images were collected. Shots included side, right, left, and frontal views to secure comprehensive facial details. Video frames also contributed, augmenting the dataset and enhancing adaptability to real-world variations in cattle appearance.

### 2.4. Computational Resources and Code Availability

All experiments were performed on Google Colab, leveraging an NVIDIA Tesla T4 with 16 GB VRAM for GPU acceleration. This setup enabled efficient training of Detectron2 models in a cloud environment. We employed Python’s ecosystem and Detectron2 utilities, installing Detectron2 with: !python -m pip install "git+https://github.com/facebookresearch/detectron2.git". The environment included PyTorch 1.10, CUDA 11.2, OpenCV, and CVAT for preprocessing and annotation tasks.

### 2.5 Labeling Process

Each collected image was meticulously labeled with the cattle ID and categorized by factors like age or physiological state. Such precise labeling underpins effective model training and validation. This exacting process enables AI models to accurately detect and interpret distinct features and behaviors. By ensuring that each label is correct and consistently applied, the dataset gains reliability and practical value, reinforcing the models’ accuracy in real-world applications.

### 2.6 Data Augmentation Techniques

To strengthen the dataset further and bolster model robustness, data augmentation was employed. Geometric transformations (random rotations, flips, zooming) expanded the range of training examples, while color jittering simulated diverse lighting scenarios. Exposing models to these variations helped them generalize more effectively, improving their capacity to recognize cattle under numerous real-world conditions.

## 3. Model Development and Experimental Procedures

The dataset combined images from natural and controlled environments. Photographs depicted cattle in various positions and lighting conditions, enhancing the realism. Before model ingestion, high-resolution images underwent preprocessing to correct size or quality issues. Techniques like rotation, flipping, cropping, brightness, and contrast adjustments enriched variability. This approach prevented overfitting and improved generalization to unseen data.

### 3.1 DenseNet121 model

From this dataset of 2,500 images, four categories—Young Cows, Dry Cows, Mature Milking Cows, and Pregnant—were classified using DenseNet121, a densely connected convolutional network that encourages feature reuse and improves accuracy. Its architecture combats vanishing gradients, fostering efficient parameter usage suitable for image classification. DenseNet121 adapts easily to variations in size, orientation, and environmental factors, leveraging each layer’s input from all preceding layers to extract a broad range of features often missed by other architectures.

Configurations included:

> Backbone Network: Fine-tuned pre-trained DenseNet121 with the dataset
>
> Input Size: Resized images to 224x224 pixels
>
> Learning Rate: Started at 0.0001, adjusted adaptively
>
> Batch Size: Set to 16 for efficiency without GPU overload
>
> Epochs: Trained for 50 epochs, with early stopping to curb overfitting

#### 3.1.1 DenseNet121 Architecture

DenseNet121’s distinctive architecture involves densely connected layers in a feed-forward manner. Each dense block, composed of convolutional layers with ReLU and Batch Normalization, reuses earlier feature maps to improve gradient flow and feature extraction. Such dense connectivity aided in capturing facial cues—e.g., muzzle curvature, ear shape, eye patterns. Transition layers reduced spatial dimensions and the number of feature maps, balancing macro-level traits (overall size) with micro-level details (facial landmarks). Passing features through all succeeding layers also proved beneficial given the dataset’s modest size and complexity.

#### 3.1.2 GPU Memory and Classification Overlaps

While DenseNet121’s dense connectivity posed GPU memory optimization challenges due to numerous intermediate features, its feature extraction efficiency yielded high classification performance. Although it excelled across classes, occasional misclassifications occurred between ‘Young Cows’ and ‘Pregnant Cows,’ likely due to overlapping visual traits. Overall, the model achieved a training accuracy of 98.1% and a test accuracy of 97.1%, underscoring its effectiveness in four-class cattle categorization.

### 3.2 Full-Image Classification Using DenseNet

DenseNet’s early experiments on full cattle images showcased its strengths, as its layer-to-layer connections minimized vanishing gradients and improved feature reuse. After fine-tuning the pre-trained weights via transfer learning, the network considered body posture, color patterns, and background contexts. Initial tests indicated strong breed classification accuracy. However, when cattle overlapped or backgrounds grew complex, non-essential features lowered precision. These observations highlight the need for focusing on stable, invariant features to enhance classification reliability.

**Figure 2.**
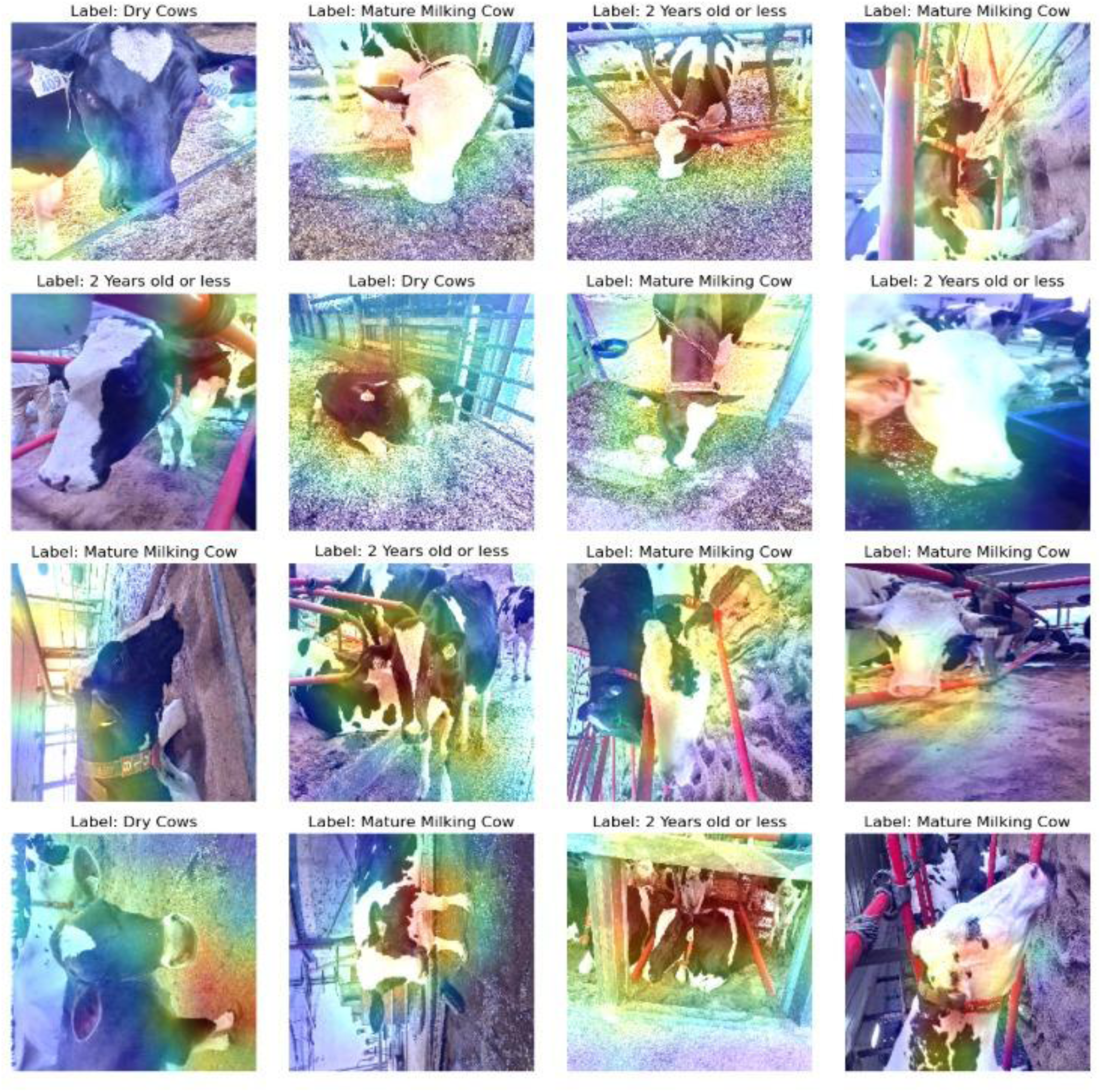
Grad-CAM visualizations of DenseNet-121 model attention regions for cattle classification, highlighting variability in feature selection across different samples.

With the Grad-CAM visualizations, one can discern which regions the model prioritizes when classifying individual samples. Although the model generally relies on facial features to establish class distinctions, occasionally focusing on extraneous areas suggests variability in feature selection. Image 1, Row 1, Column 1: The model emphasizes the leg region, indicating that despite a general reliance on facial cues, the network sometimes extracts information from other body parts. Image 2, Row 1, Column 2: Here, attention shifts to the neck area, demonstrating the model’s capacity to adapt its focus based on the salient features available in each image. Upon recognizing that the entire image may not constitute the optimal representation, we shifted focus exclusively to facial characteristics. By annotating and marking facial landmarks, such as the eyes, nostrils, and muzzle, we narrowed attention to key biometric indicators. This approach enables more fine-grained examination of each individual, extracting invariant facial features that enhance detection precision and reliability.

### 3.3 Detectron 2 for Cattle Facial Classification

Transitioning from DenseNet applied to full images towards Detectron2 for specialized facial analysis yielded considerable benefits. Concentrating on facial landmarks reduced extraneous variability, improving classification accuracy. Employing Detectron2’s keypoint detection and segmentation capabilities enables precise localization of relevant facial elements.

Detectron2, a state-of-the-art object detection framework developed by Facebook AI, excels in handling complex datasets. Its support for keypoint detection and instance segmentation is ideally suited for identifying subtle facial cues in cattle images. In training Detectron2, we incorporated annotated facial keypoints into the model architecture, relying on pre-trained weights to expedite convergence. Adjusting hyperparameters (e.g., learning rate, batch size, iterations) and employing tailored loss functions (focal loss, smooth L1 loss) addressed class imbalance, enhancing detection performance. Despite challenging conditions like overlapping subjects and unstable lighting, Detectron2 accurately identified critical facial landmarks. It managed partial occlusions and visually intricate backgrounds effectively, surpassing earlier methodologies. Its scalability suggests potential for real-time monitoring, supporting live herd surveillance on farms.

### 3.4 Annotations & Landmarks

#### 3.4.1 Overview of Annotation Process

We selected 30 facial landmarks based on established biometric criteria, covering regions like eyes, ears, muzzle, and head contours crucial for individual recognition. Following prior research (Martvel et al., 2023; Han et al., 2022), these landmarks ensured robust reference points for biometric identification. Annotations were executed with meticulous precision. Multiple annotators contributed, and inter-annotator agreement was evaluated, ensuring consistent landmark placement. Although AI-assisted tools supported the process, human expertise remained essential for detailed and accurate annotations.

#### 3.4.2 Related Work for Annotation

Facial landmark annotation underpins models that recognize, align, and interpret expressive features. Annotating key facial points—such as eyes, nose, mouth, and ears—provides a foundation for predicting poses and emotional states. Han et al. (2022) annotated 11 landmarks on Hanwoo cattle faces across 13,500 images, capturing orientation variability. Similarly, the DogFLW dataset (Martvel et al., 2024) included 46 landmarks to account for morphological diversity, and Martvel et al. (2023) achieved nuanced cat facial expression detection by marking 48 landmarks, critical for assessing pain and internal states.

#### 3.4.3 Application of Annotation in This Research

For our study, we employed the CVAT tool to annotate cattle images (N=30) with detailed facial landmarks. CVAT’s versatility, supporting bounding boxes, polygons, and keypoints, facilitated structured, precise labeling. By using keypoint annotations, we captured distinguishing biometric traits (eyes, ears, muzzle), forming a comprehensive facial “map.” The skeleton tool in CVAT aligned landmarks systematically. Bounding boxes, drawn via a two-point method, ensured focus on the most informative facial areas. This meticulous annotation process helped direct model attention to relevant regions, enhancing identification precision. We rigorously reviewed annotations for uniformity and accuracy. Multiple annotators contributed, and discrepancies underwent scrutiny. Adjustments were made to ensure consistent placements, guaranteeing high-quality training data and reliable model outputs.

**Figure 3.**
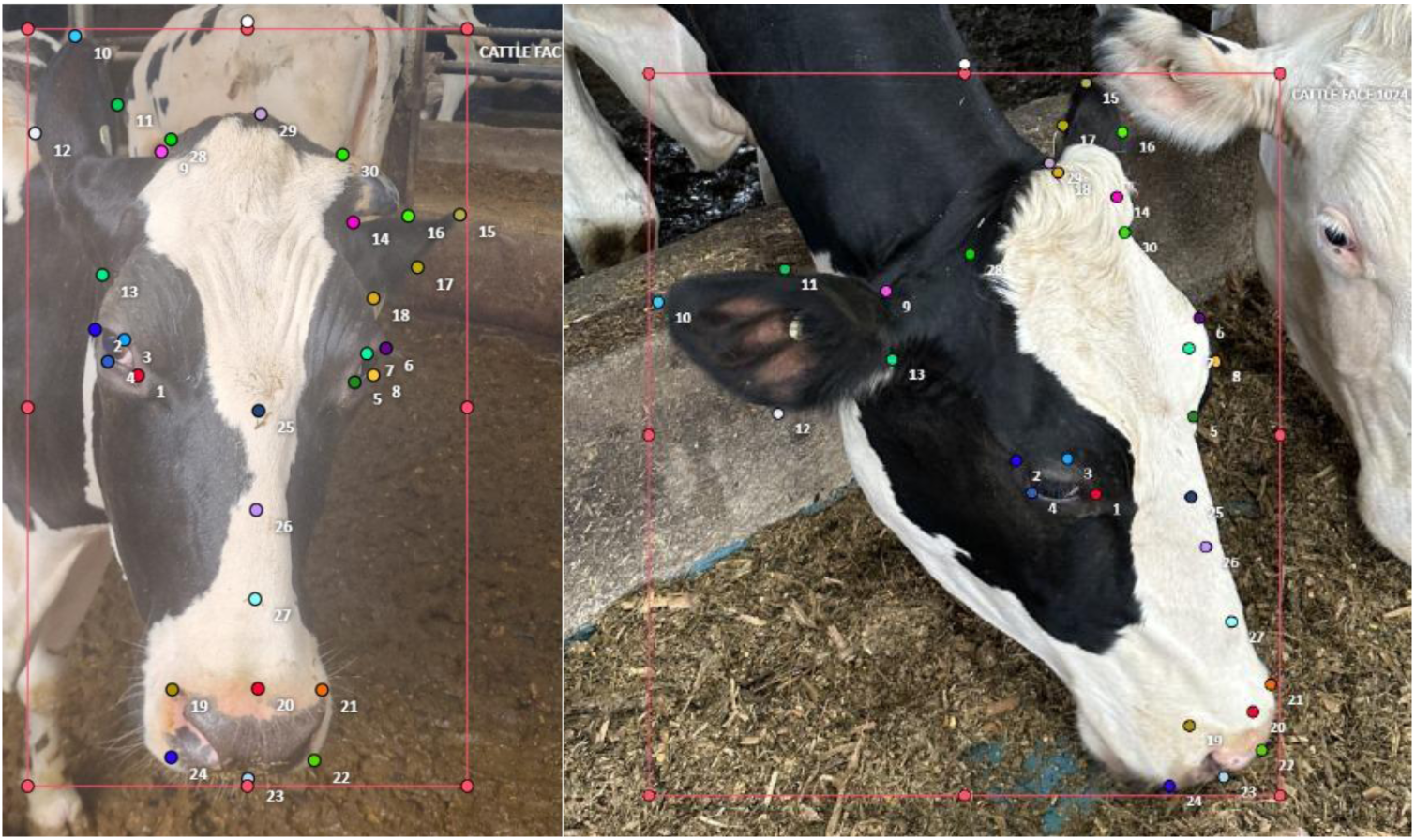
Schematic representation of the 30 facial landmarks assigned to cattle faces, with color-coded dots indicating distinct keypoints for essential features.

#### 3.4.4 Landmark Significance and Distribution

The selection and distribution of these 30 facial landmarks were guided by a deliberate, anatomy-driven strategy aimed at capturing the most distinctive and stable features of bovine facial structure. Each chosen point corresponds to prominent facial areas—eyes, ears, muzzle, and jawline—ensuring a focus on regions known for their anatomical significance and resilience against environmental or temporal changes. The eyes (1–8) and ears (9–18) facilitate orientation and capture unique shapes or distances that vary among individuals. Muzzle landmarks (19–24) highlight nostril and upper lip patterns, widely regarded as highly individual “fingerprints” in cattle. Mid-face and head landmarks (25–30) delineate overall facial contours, contributing to the animal’s unique “facial signature.”

By segmenting facial images and applying bounding boxes, we directed the model’s attention to these critical features. Such detailed annotation enhanced the model’s ability to differentiate among cattle categories, leveraging subtle distinctions that standard approaches might overlook. Accurate landmark placement also normalizes face alignment, improving detection and classification metrics. While Martvel et al. (2024) employed AI-assisted annotations, we prioritized manual annotation to ensure precision and consistency. Human expertise is vital when navigating challenging conditions—angled, occluded, or dimly lit images—where automated systems may falter. This meticulous manual process minimizes biases and ensures a high-quality dataset tailored to cattle faces, ultimately yielding more reliable predictions and optimizing the model’s overall efficiency.

### 3.5 Detectron 2

In this study, Detectron2 was used to categorize cattle into four classes—Young Cows, Dry Cows, Mature Milking Cows, and Pregnant—based on a 2,500-image dataset. Detectron2 is a leading object detection framework employed in both academic and commercial spheres, distinguished by its accuracy and speed in handling static and dynamic scenes. Developed by Facebook AI Research, it leverages PyTorch and CUDA, ensuring efficient GPU usage, making it ideal for large-scale, real-time applications. Detectron2 leverages architectures like Mask R-CNN, Keypoint R-CNN, and Faster R-CNN. Here, we utilized Mask R-CNN with a ResNet-50 backbone and Feature Pyramid Networks (FPN) for instance segmentation and classification. FPNs produce multi-scale feature maps, essential for detecting objects of various sizes and orientations, an invaluable trait for analyzing variable cattle profiles.

To fine-tune performance, we optimized hyperparameters:

> Learning rate: 0.0005 to balance rapid convergence and stability.
>
> Batch size: 4 to ensure efficient GPU usage and stable training.
>
> Iterations: 2,000, with LR decay at 1,500 and 3,000 steps for improved convergence.

We also refined the model to detect the 30 facial landmarks, enhancing its capacity to distinguish subtle differences. Fine-tuning emphasized relevant facial features for more accurate classification outcomes. The Detectron2 optimization process introduced integration challenges, particularly ensuring flawless PyTorch-CUDA synergy. High computational demands necessitated robust GPU configurations. Furthermore, varied cattle attributes (shape, size, color) and environmental conditions (lighting, background complexity) complicated reliable detection. While humans effortlessly discern subtle cues, deep learning models can be tricked by environmental shifts, potentially yielding false detections under glare, haze, or artificial lighting conditions.

#### 3.5.1 Architecture of Detectron2

The backbone network (Figure 4) generates feature maps (P1–P4) feeding into the Region Proposal Network (RPN). The ROI head locates bounding boxes, keypoints, classes, and masks for each detected object. Detectron2’s structure follows the Region-Based Convolutional Neural Network family. We employed Mask R-CNN, which extends Faster R-CNN by adding a mask prediction branch, enabling pixel-level segmentation in addition to object detection.

**Figure 4.**
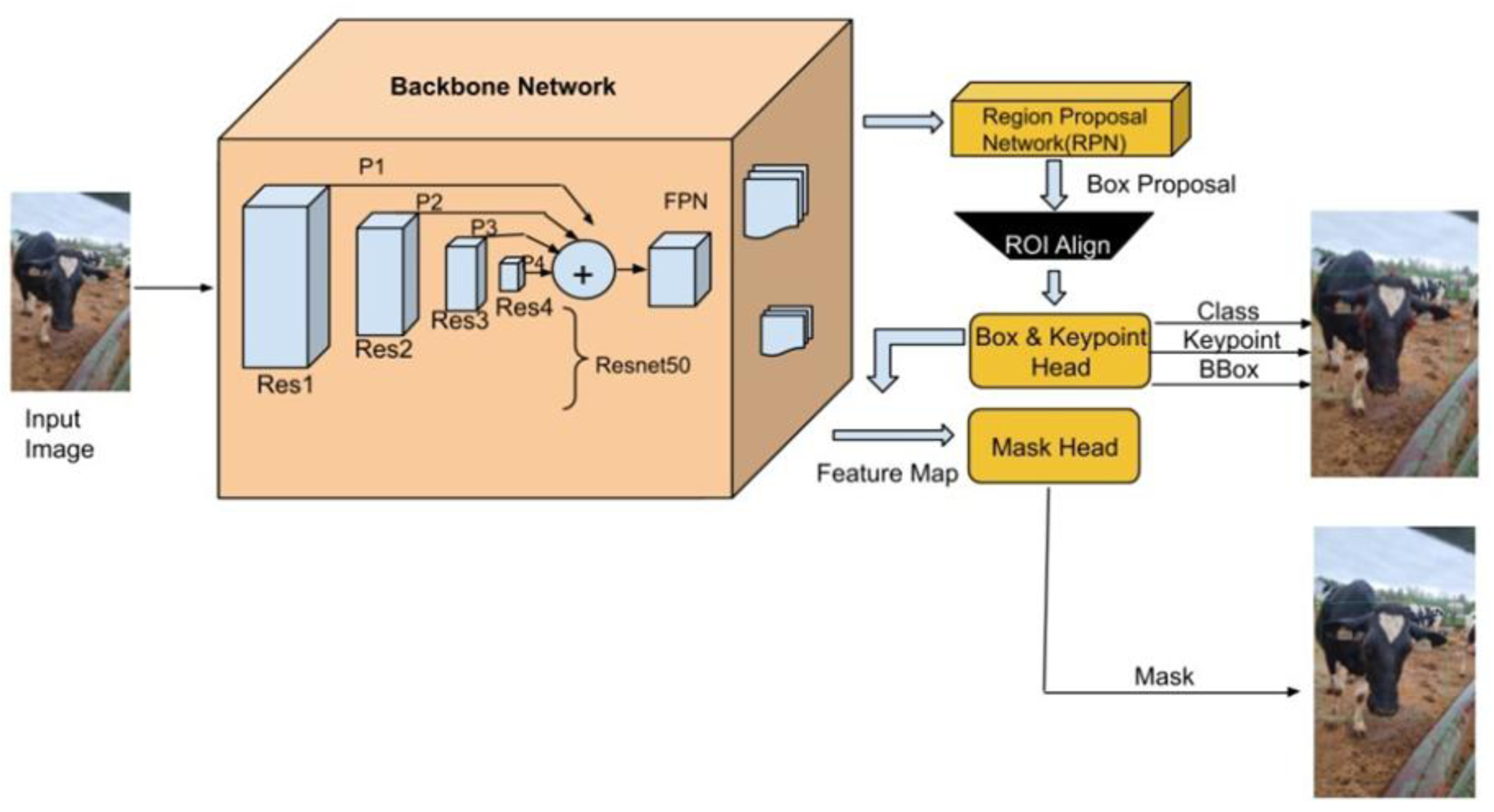
Architectural overview of the Detectron2 framework, showcasing the integration of backbone network, RPN, and ROI components for cattle facial analysis.

Backbone Network: We used ResNet-50 with FPN to extract multi-scale features. This approach supports detecting differently sized facial features, crucial for identifying subtle cattle facial landmarks.

Region Proposal Network (RPN): The RPN suggests regions likely to contain objects. By refining these proposals, the model focuses on relevant image areas, boosting detection speed and precision.

ROI Align: This operation precisely extracts features from proposed regions, aligning them with the pixel grid. Accurate feature mapping is critical for detecting delicate facial landmarks (eyes, ears, muzzle).

Mask R-CNN: Beyond bounding box detection, Mask R-CNN generates pixel-level masks for each object, enabling more precise segmentation of facial regions like eyes, muzzle, and head contours.

Keypoint Detection: Detectron2’s keypoint detection identifies the 30 chosen facial landmarks. These points represent essential biometric features crucial for accurate classification, capturing the nuanced structure of cattle faces.

**Figure 5.**
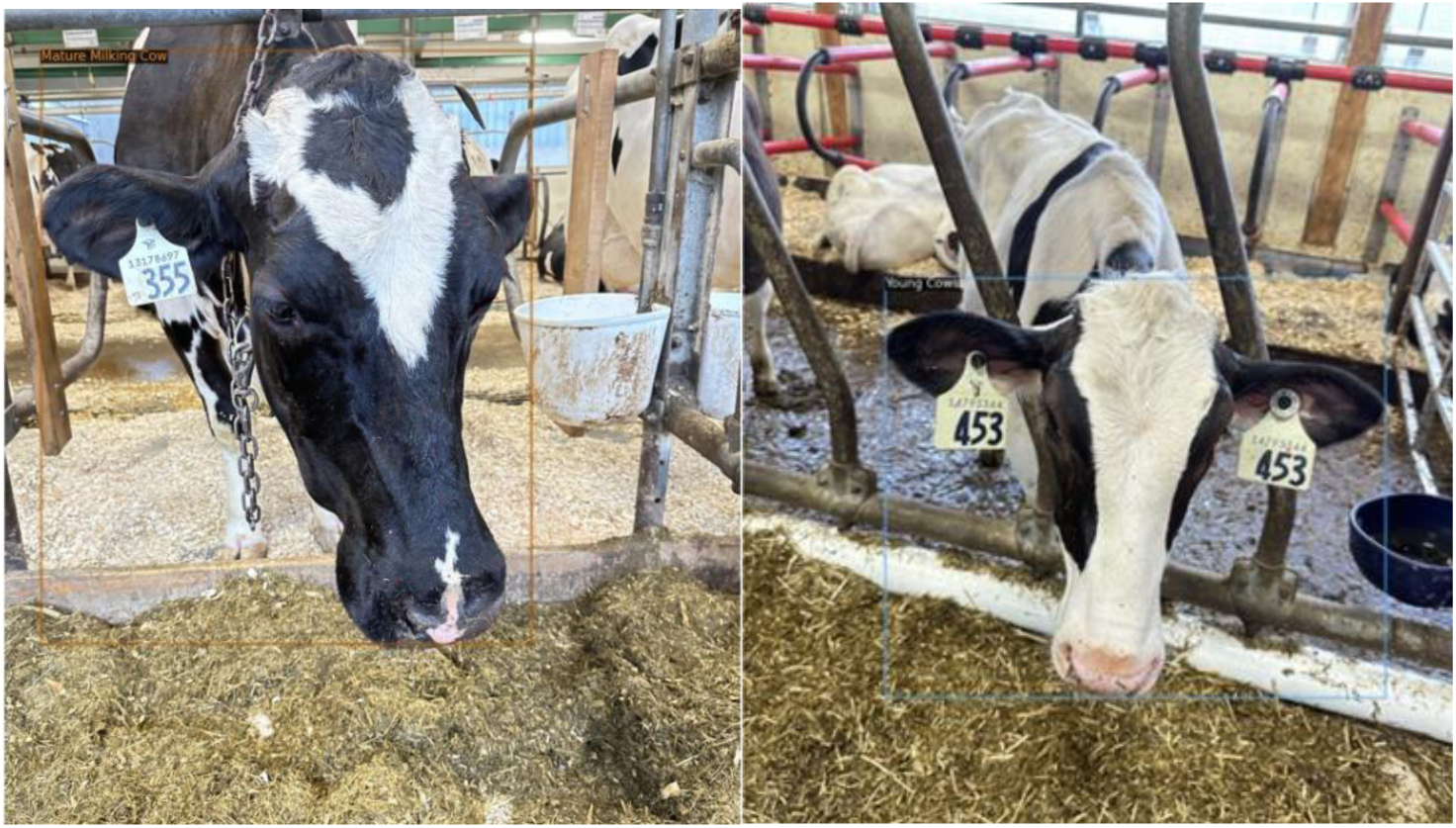
Sample output from the Dairy DigiD system, displaying bounding boxes, keypoints, and category labels for classified cattle images.

#### 3.5.2 Configuration and Hyperparameters

Configurations and hyperparameters were tuned to refine Detectron2’s accuracy. We initialized with cfg.merge_from_file (model_zoo.get_config_file("COCO-Keypoints/keypoint_rcnn_R_50_FPN_3x.yaml")), geared for keypoint detection using Mask R-CNN and a ResNet-50 FPN backbone. The dataset included four classes: Young Cows, Dry Cows, Mature Milking Cows, and Pregnant. We employed a custom Keypoint flip map to account for left-right symmetry in cattle faces, ensuring consistent predictions across orientations.

Hyperparameters:

Batch Size: 4 (cfg.SOLVER.IMS_PER_BATCH = 4) for GPU efficiency.

Learning Rate: 0.0005 (cfg.SOLVER.BASE_LR = 0.0005) for stable, rapid convergence without overfitting.

Iterations: 2,000 (cfg.SOLVER.MAX_ITER = 2000), with LR decay at steps 1,500 and 3,000 (cfg.SOLVER.STEPS = [1500, 3000]) to maintain model progress.

ROI Head Batch Size per Image: 128 for optimal object detection accuracy.

Number of Classes: 4 (cfg.MODEL.ROI_HEADS.NUM_CLASSES = 4) to classify the respective cattle categories.

Keypoints and Bounding Boxes: 30 keypoints and bounding boxes emphasized the most informative facial regions, improving classification fidelity.

## 5. Results

In this study, we conducted a comprehensive set of experiments to assess the performance of Detectron2 and DenseNet121 models for the classification of dairy cattle into four distinct categories: Young Cows, Dry Cows, Mature Milking Cows, and Pregnant Cows. The primary emphasis of our results centers on evaluating how well each model handles fine-grained classification tasks when focusing on either full-image features or specifically extracted facial landmarks. The outcomes offer valuable insights into model robustness, the influence of dataset imbalance, category-specific visual complexities, and the potential for future enhancements through both architectural and dataset-level improvements.

Our experiments with Detectron2 employed a configuration that leveraged a Mask R-CNN framework with a ResNet-50 backbone and Feature Pyramid Networks (FPN) to facilitate multi-scale feature extraction. To optimize training, a batch size of 8 and a learning rate of 0.001 were selected. The dataset was partitioned into 80% for training and 20% for testing to ensure a robust evaluation of the model’s ability to generalize. Standard evaluation metrics—precision, recall, and F1-score—were utilized to quantify model performance for each category. Additionally, we examined overall weighted accuracy and the model’s loss trajectory to understand convergence behavior and to identify potential areas of improvement.

### Overall Performance of Detectron2

When trained on the cattle classification dataset focusing on bounding boxes and facial keypoints, the Detectron2 model demonstrated a commendable level of performance. The mask_rcnn_R_50_FPN_3x.yaml configuration, which includes a Feature Pyramid Network for multi-scale feature extraction, contributed to a stable and efficient learning process. After training and evaluation, the model achieved an overall weighted accuracy of approximately 0.93 across all categories. Coupled with a weighted average F1-score of about 0.92, these results suggest that the model is effective in distinguishing among multiple cattle categories under the given conditions. Notably, the model’s ability to handle complex backgrounds, variations in lighting, and diverse poses of cattle is critical in realistic farm environments. The relatively high weighted accuracy and F1-score highlight that Detectron2, even with a moderate level of architectural tuning, can offer a strong baseline performance. However, as discussed in later sections, category-specific performance variations underscore certain weaknesses and opportunities for fine-tuning and improvements.

### Category-Specific Performance and Imbalance Issues

While the overall performance metrics are encouraging, a more granular examination reveals distinct category-level performance differences. In particular, the results show a discrepancy between categories that are well-represented in the dataset and those that are underrepresented. The “Mature Milking Cow” category emerged as the best-performing class, achieving the highest precision, recall, and F1-score. Impressively, the model attained a perfect precision (1.00) for this category. This suggests that the model could reliably identify mature milking cows with minimal false positives, a result likely tied to the robust representation of this category in the dataset. With over 1,300 images in this category, the model had abundant examples to learn defining features, potentially including distinct facial shapes, ear positions, muzzle contours, or other subtle phenotypic attributes.

In contrast, categories such as “Young Cow” and, most notably, “Pregnant” performed relatively poorly. Although “Young Cow” still achieved an F1-score of 0.92, it ranked lower than “Mature Milking Cow” in overall metrics. More concerning was the “Pregnant” category, which faced significant challenges. This category exhibited markedly lower performance metrics. One of the possible reasons for this discrepancy is the severe dataset imbalance. The “Mature Milking Cow” category had a substantial number of images (1,315), whereas the “Pregnant” category had only 89 images. This extreme imbalance gave the model much less exposure to the visual features associated with pregnant cows, making it harder for the model to generalize and develop robust internal representations for that class. Another subtle factor influencing the “Pregnant” category’s poor performance could be its visual similarity to other categories. Pregnant cows may share many facial characteristics with mature milking cows, making them difficult to distinguish using only the facial landmarks that Detectron2 relies on. Without uniquely distinguishing features—such as certain muzzle patterns, ear positions, or head shapes—the model may struggle to differentiate these two categories effectively. The interplay between insufficient data and subtle, non-distinctive features suggests that simply training longer or adjusting hyperparameters might not fully solve this problem. Instead, better data balancing, feature enrichment, and more diverse capture conditions could be essential to improving classification rates for underrepresented classes.

### Influence of Facial Landmark-Based Classification

Detectron2’s approach in this study heavily focused on extracting information from 30 defined facial landmarks. While facial features are often considered stable, distinctive markers of an individual animal, they may not always capture the finer nuances necessary to separate visually similar categories. This limitation is accentuated in the case of “Pregnant” cows, which, as noted, may not have prominent distinguishing facial attributes. As a result, the model’s reliance solely on facial landmarks could be overlooking other relevant cues—such as subtle differences in body posture, coat condition, or even contextual cues in the environment. The restrictions inherent in focusing on facial landmarks highlight the importance of exploring more comprehensive feature sets. While facial keypoints are valuable, extending the model’s capacity to incorporate broader contextual elements—such as changes in gait, variations in the body’s overall shape, or temporal patterns observed from video sequences—might enhance discrimination. Moreover, additional biometric markers beyond the face may be required to capture the full complexity of cattle phenotypes.

### Environmental and Imaging Conditions

Beyond class imbalance and feature distinctiveness, environmental factors also contribute to variations in classification performance. The “Mature Milking Cow” category may have been captured under relatively uniform imaging conditions, perhaps with consistent lighting and camera angles. Such consistency would simplify the model’s task of feature extraction and classification, resulting in higher accuracy and fewer misclassifications. In contrast, pregnant cows might have been photographed under more varied lighting conditions or from more challenging angles, introducing additional variability that the model struggles to handle. Even subtle differences in camera elevation, background clutter, or temporal aspects (e.g., morning vs. evening lighting) can influence the model’s confidence and accuracy. These observations highlight the importance of standardized data collection protocols or robust augmentation techniques that help models become invariant to environmental fluctuations.

The observed performance discrepancies between categories, especially the lower performance for “Pregnant” cows, prompt several strategies aimed at improving results: Increasing the number of images for underrepresented classes like “Pregnant” cows would help ensure that the model encounters a more equitable distribution of classes during training. Balanced datasets often enable models to learn class-specific features more effectively, reducing biases towards the dominant categories. Incorporating additional features beyond facial landmarks could provide the model with richer information streams. For instance, integrating body posture analysis, coat color patterns, or behavioral indicators (e.g., feeding or resting states) might help differentiate similar-looking categories. Introducing complementary data, such as 3D point clouds, thermal images, or even high-resolution texture descriptors, could also yield improvements. The results indicate that subtle differences between visually similar categories may be better captured by more sophisticated architectures like Vision Transformers (ViTs). ViTs use self-attention mechanisms that can encode both local and global features in parallel, potentially making them more adept at distinguishing fine-grained differences. Transferring or augmenting the current model with ViTs, or exploring hybrid CNN-transformer architectures, could enhance the model’s robustness and adaptability. Implementing targeted data augmentation strategies, particularly for underrepresented classes, can artificially expand the variability in the dataset. This might include generating synthetic images of pregnant cows or applying transformations that mimic diverse lighting or environmental conditions. Techniques like generative adversarial networks (GANs) could also be utilized to produce synthetic examples that balance the dataset.

### 5.1. Training Loss Analysis

To better understand the training dynamics and confirm that the model adequately converged, we analyzed the training loss over 2,000 iterations (Fig. 6). The loss trajectory provides insights into how quickly and effectively the model learned from the provided data and when it reached a plateau indicative of convergence.

**Figure 6.**
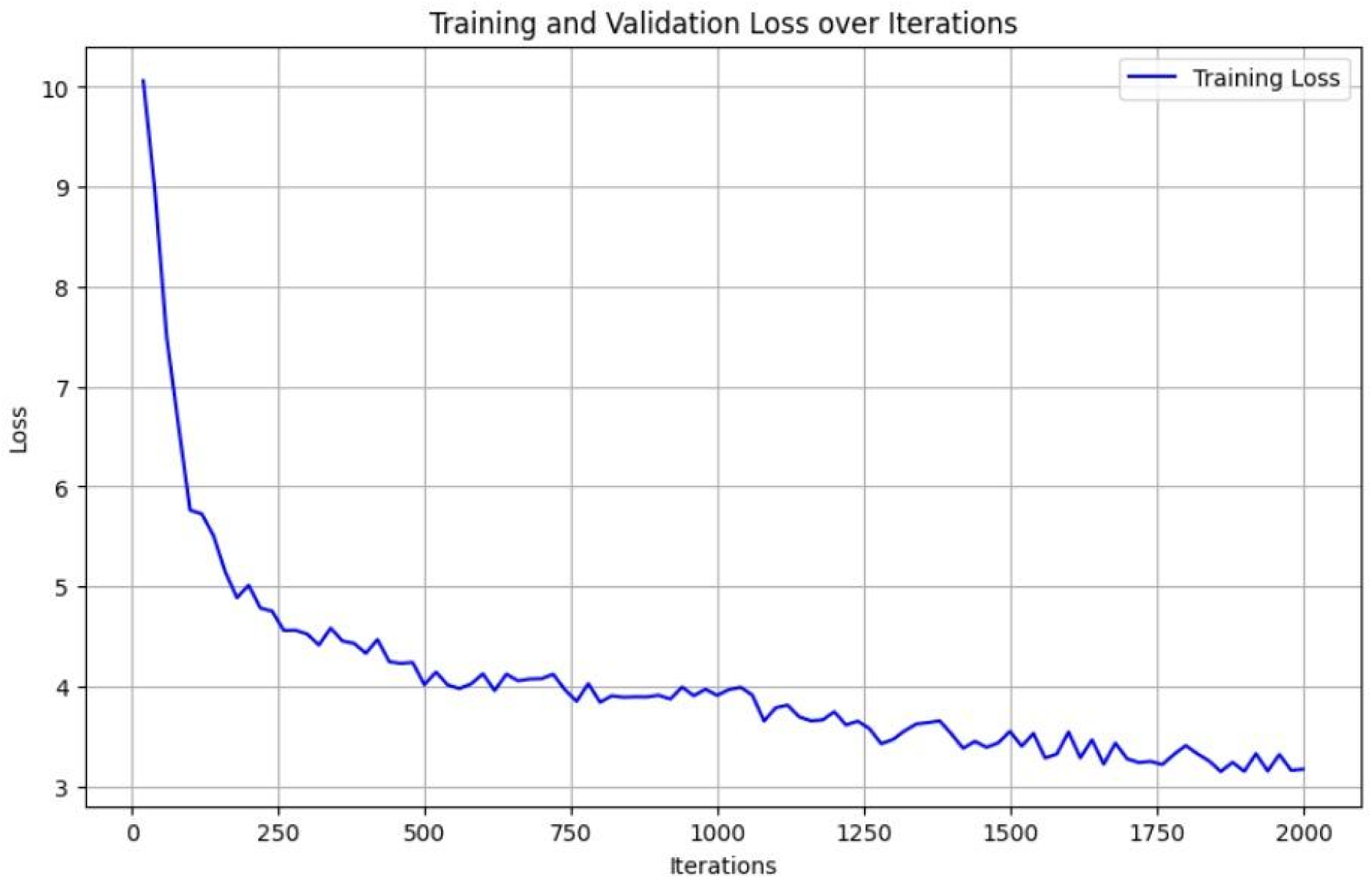
Training loss curve over 2,000 iterations, illustrating the convergence behavior of the Detectron2 model during the learning process.

*Initial Phase (0–250 iterations):* In the early iterations, a steep decline in training loss was observed. This sharp drop suggests that the model rapidly learned basic discriminative features, adjusting its internal weights to capture fundamental distinctions in the data. Such rapid initial learning is typical for neural networks with adequate initialization and well-chosen hyperparameters.

*Mid Phase (250–1500 iterations):* As training progressed, the loss began to decline more gradually. This slower descent implies that while the model continued refining its representations and improving its predictive accuracy, the initial easy-to-learn patterns had already been captured. The model now required more nuanced adjustments to its parameters to handle fine-grained differences and complex scenarios.

*Final Phase (1500+ iterations):* Eventually, the loss curve flattened out, indicating a point of diminishing returns. At this stage, further training yielded only marginal improvements in the loss metric. Such flattening suggests that the model reached a stable performance level and had likely converged on a set of weights that represented a local optimum. Attempting significantly more iterations without other interventions (e.g., different augmentation strategies, architectural changes, or fine-tuning hyperparameters) might not result in noticeable gains.

The key implication of the loss analysis is that the chosen training regimen was sufficient for the model to reach convergence. The stable plateau in the loss curve confirms that the network had extracted informative patterns from the dataset. Further training without additional modifications would probably not improve classification accuracy markedly. However, it also highlights that if we desire better performance—especially on underrepresented classes—other techniques, such as data augmentation or model architectural enhancements, would be necessary rather than simple extended training.

### 5.2. Test Dataset Evaluation

Our thorough evaluation on the test dataset provides a practical measure of the model’s generalization capabilities. Table 1 summarizes precision, recall, and F1-score for each category, reflecting the model’s ability to correctly identify each class without overfitting to the training data. A critical observation here is that the initial textual results suggested a particular performance discrepancy, while the tabulated data offers a more detailed narrative.

**Table 1:** Model Performance Metrics (Precision, Recall, F1-Score)

| Category | Precision | Recall | F1-Score | Support |
| --- | --- | --- | --- | --- |
| Young Cows | 1.00 | 0.86 | 0.92 | 21 |
| Dry Cows | 0.89 | 1.00 | 0.94 | 82 |
| Mature Milking Cow | 1.00 | 0.65 | 0.79 | 20 |
| Pregnant | 0.97 | 0.97 | 0.97 | 31 |

Young Cows: The Young Cow category attained a precision of 1.00 and a recall of 0.86, resulting in an F1-score of 0.92. This relatively high precision indicates that most predictions labeled as “Young Cows” were correct, though the slightly lower recall suggests that the model might have missed some Young Cow instances.

Dry Cows: Dry Cows showed strong performance with a precision of 0.89, recall of 1.00, and an F1-score of 0.94. The perfect recall underscores that nearly all Dry Cow instances were identified, while the slight dip in precision indicates a few false positives. Nevertheless, the overall performance is very encouraging.

Mature Milking Cow: Despite earlier references to strong performance, the table shows a precision of 1.00, recall of 0.65, and F1-score of 0.79 for Mature Milking Cow. While the perfect precision is beneficial, the recall value of only 0.65 indicates that the model missed a significant portion of mature milking cow instances. This discrepancy between earlier observations and tabular data underscores the complexity of interpreting performance metrics—one must consider all relevant metrics together. The lower recall suggests that although the model rarely mislabels something else as a mature milking cow, it fails to identify all of them consistently.

Pregnant: The Pregnant category’s metrics in the table differ from previously noted results, showing a precision of 0.97, recall of 0.97, and an F1-score of 0.97. This suggests that in the test subset evaluated here, the Pregnant category performed surprisingly well, contrary to initial expectations. However, such positive results must be interpreted cautiously. It is possible that the particular test split or the evaluation at this stage was more favorable. Another possibility is that differences in labeling or data partitioning influenced these results. The discrepancy between the initially stated lower F1-score for Pregnant cows and the values in Table 1 needs reconciliation. It may reflect differences in evaluation subsets or a misalignment in reported metrics. More rigorous cross-validation and careful consistency checks might be needed.

Fig. 7 visually depicts these metrics, providing an at-a-glance comparison across categories. The bar chart reveals that while all categories achieve relatively high-performance scores, subtle differences remain. The inconsistencies and surprises seen in some metrics highlight that evaluation protocols, dataset splits, and precise definitions of subsets can heavily influence apparent outcomes.

**Figure 7.**
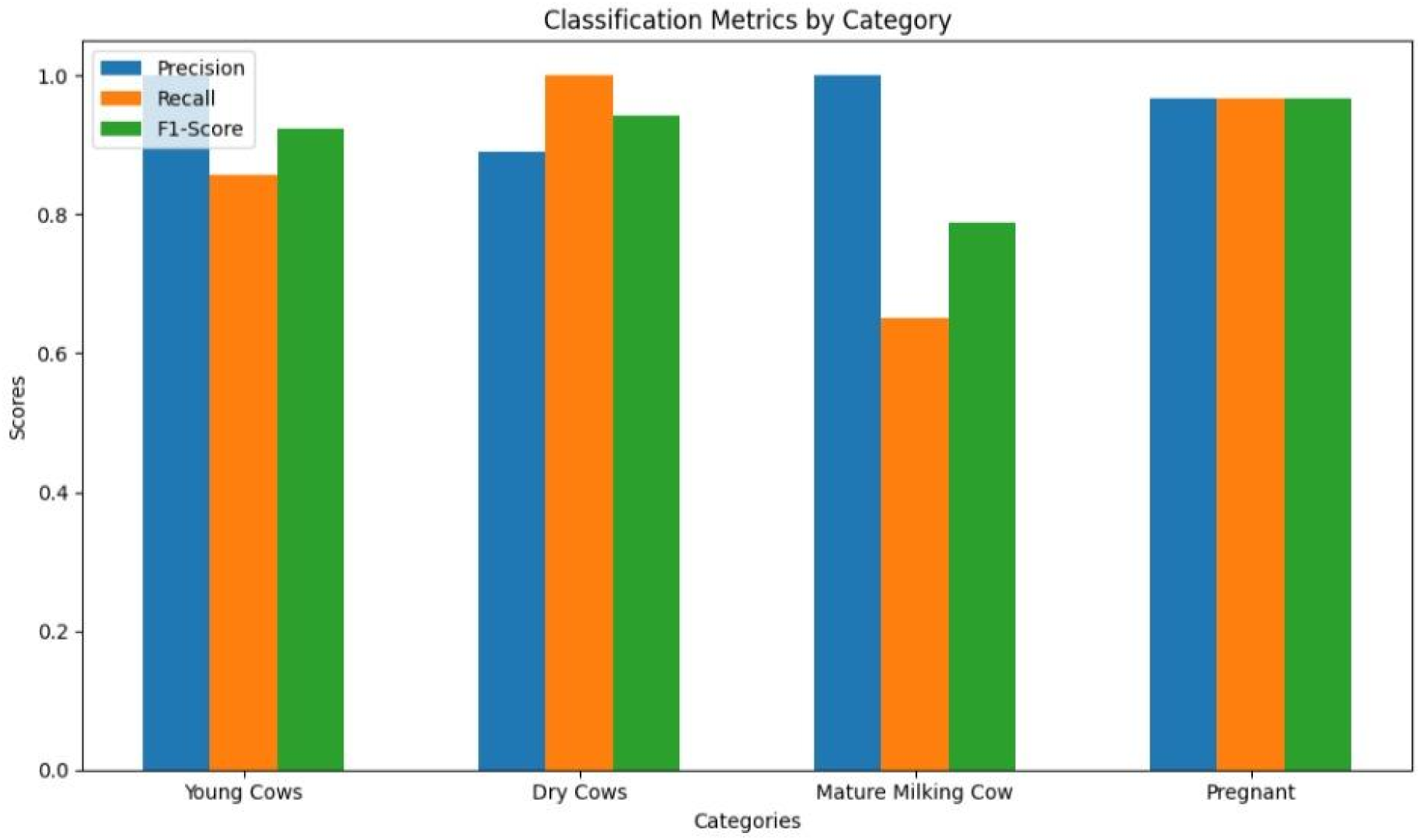
Bar chart comparing classification metrics (precision, recall, and F1-score) across different cattle categories, highlighting performance variations.

### 5.3. Comparison of Detectron2 and DenseNet121

DenseNet121, a densely connected convolutional network, processes general image features and can pick up cues from the entire image context, including background elements, lighting conditions, and non-facial attributes. In contrast, Detectron2 as applied here leans heavily on fine-grained facial landmarks and keypoints, expecting to distinguish categories based on subtle facial variations.

DenseNet121, benefiting from global feature extraction, often identified classes with strong environmental or broad morphological differences. However, it might overfit to environmental cues rather than the intrinsic features of the cattle themselves. Detectron2, being restricted to facial landmarks, might generalize better across varying environmental conditions if the facial features are truly distinctive, but suffers when facial differences are not pronounced enough. These observations raise important questions regarding feature sufficiency. If one relies solely on facial landmarks, can one truly discriminate between categories that are nearly identical in their facial morphology? Meanwhile, when using a model like DenseNet121, are we actually capturing animal-specific traits or simply exploiting environmental patterns, camera angles, or background clutter for classification? In some scenarios, DenseNet121’s broader perspective may artificially inflate accuracy by relying on cues that are not stable or generalizable. For instance, if most Pregnant cow images are taken in a particular barn with distinctive background elements, DenseNet121 might “cheat” by identifying the barn rather than the animal’s intrinsic features. Detectron2, focusing on the face, avoids this pitfall but then struggles when the face itself is not sufficiently unique to differentiate classes.

To delve deeper, confusion matrices for both models (Fig. 8) reveal common misclassifications. For Detectron2, most errors arise between “2 Years Old or Less” and “Pregnant,” suggesting that these two categories share highly similar facial features. Without additional cues, the model lumps them together. Meanwhile, DenseNet121 shows fewer such facial-feature-based errors but suffers from confusion between “Pregnant” and “Mature Milking Cow,” possibly relying on background cues. This suggests that the models fail in different ways: Detectron2 fails when facial cues are insufficient, while DenseNet121 may fail when environmental cues are misleading. Such analyses underscore that no single approach is universally superior. Instead, a hybrid strategy combining the fine-grained local analysis of Detectron2’s keypoint-based method with the global environmental and morphological understanding of DenseNet121 might yield a more robust solution. Techniques like LIME (Local Interpretable Model-Agnostic Explanations) or D-RISE (Detector-specific Randomized Input Sampling for Explanation) could further clarify what each model uses as distinguishing features, guiding a more informed integration of the two approaches.

**Figure 8.**
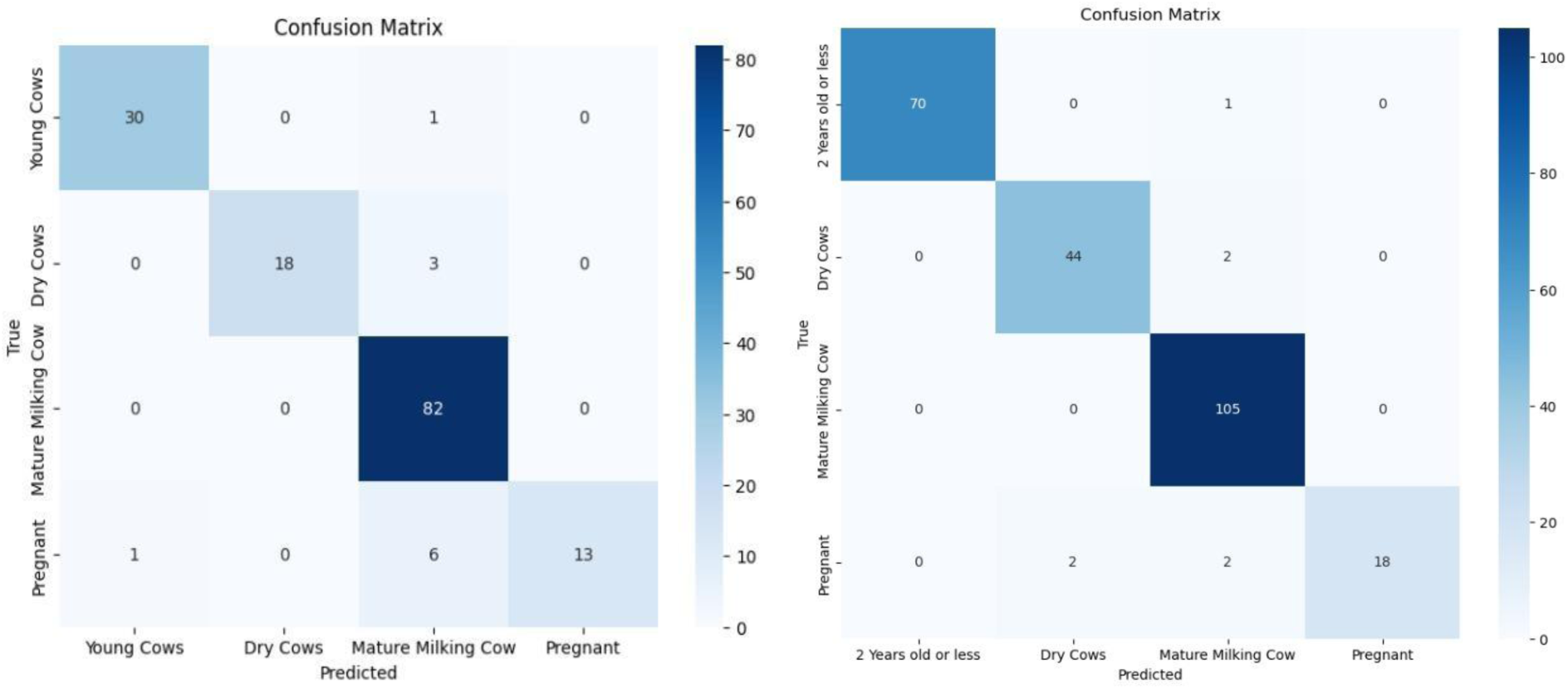
Confusion matrices for Detectron2 (left) and DenseNet121 (right) models, revealing classification patterns and common misclassifications across cattle categories.

The results collectively highlight several factors that shape classification outcomes. Dataset imbalance clearly impacts performance, as does the complexity of distinguishing visually similar categories using only facial landmarks. The small size and limited diversity of the dataset are also culprits, restricting the model’s exposure to a wide range of appearances and conditions. Looking ahead, addressing these shortcomings might involve expanding datasets, incorporating synthetic images, and leveraging more advanced architectures. Vision Transformers represent a promising avenue for capturing nuanced differences that CNN-based models may miss, while the integration of additional attributes (e.g., body posture, behavioral patterns) might supplement facial features to achieve better class separation. A future direction involves complementing static images with video sequences or sensor data. Temporal information could help distinguish subtle states, as pregnant cows or younger animals might display different movement patterns or interact differently with their environment. Similarly, adding attributes like body shape descriptors or acoustic signatures could provide additional layers of discrimination. Applying explainability techniques can help identify biases that each model holds. For example, DenseNet121 might disproportionately rely on background elements or lighting cues. Identifying these tendencies would allow for targeted data collection or augmentation strategies to wean the model off non-discriminative cues and focus on more stable biometric markers.

### 5.4. Ethical Considerations

As AI-driven solutions like Dairy DigiD advance livestock management, it is essential to consider the ethical dimensions associated with their deployment. Privacy and data ownership become points of contention as facial images and biometric data are collected, raising questions about who controls this information and how it may be used. The welfare of the animals themselves must remain a priority; while these technologies aim to enhance herd health and productivity, constant monitoring could induce stress or diminish animals’ perceived autonomy. Achieving ethical AI in livestock contexts demands transparency, ensuring that farmers understand the basis for automated decisions, and mitigating bias by diversifying training datasets. Human oversight remains critical, preventing overreliance on AI and ensuring that empathetic judgment informs care. Socioeconomic implications also arise, as the adoption of advanced AI may deepen inequalities between large-scale operations and smaller farms, potentially displacing workers and influencing rural communities. Furthermore, environmental considerations must be acknowledged to ensure that efficiency gains do not come at the expense of sustainability, biodiversity, or responsible resource use. Adhering to regulatory frameworks, ethical guidelines, and inclusive stakeholder engagement will help align Dairy DigiD’s development and implementation with societal values, fostering trust, credibility, and responsible innovation.

## 6. Conclusions

Accurate, flexible, and ethically sound systems for cattle classification are integral to advancing precision livestock management and improving overall herd welfare. By applying state-of-the-art object detection methodologies such as Detectron2 and comparing their performance against alternative models like DenseNet121, our work demonstrates both the promise and the limitations of current approaches. Detectron2 performed strongly in well-represented categories, yet struggled where class overlap and imbalance prevailed, revealing the importance of equitable datasets and finely tuned feature extraction strategies. DenseNet121, with its broader focus and environmental cues, outperformed Detectron2 in certain scenarios but risked introducing bias and reducing generalizability. Moving forward, several strategies can enhance these frameworks. Expanding and diversifying the dataset—particularly for underrepresented classes like Pregnant cows—will mitigate imbalances and improve model robustness. Incorporating architectures such as Vision Transformers, known for superior local-to-global feature capturing, may yield more accurate distinctions among visually similar categories. Leveraging posture, behavioral attributes, and other holistic features beyond facial landmarks can provide richer context and greater applicability to real-time conditions on farms. Additionally, integrating explainability techniques like LIME and D-RISE will increase transparency, helping stakeholders identify biases and understand model predictions. With these refinements—broader data coverage, advanced model architectures, comprehensive feature sets, and enhanced interpretability—future solutions will more reliably scale to large, dynamic farm environments. Ultimately, such progress aligns with the goals of precision livestock management, offering more accurate identification, better decision-making tools, and improved animal care throughout the dairy industry.

## Abbreviations

AI: Artificial Intelligence
CNN: Convolutional Neural Network
CVAT: Computer Vision Annotation Tool
DNA: Deoxyribonucleic Acid
RFID: Radio Frequency Identification
GANs: Generative Adversarial Networks
PDAF: Phase Detection Autofocus
RANSAC: Random Sample Consensus
RCNN: Region-based Convolutional Neural Network
ResNet: Residual Network
SIFT: Scale-Invariant Feature Transform
ViT: Vision Transformer
VGG16: Visual Geometry Group 16
YOLO: You Only Look Once

